# Analysis of archaic human haplotypes suggest 5-hmC to act as epigenetic guide for meiotic point recombination

**DOI:** 10.1101/227702

**Authors:** Bernett Lee, Samantha Cyrill, Wendy Lee, Rosella Melchiotti, Anand Andiappan, Michael Poidinger, Olaf Rötzschke

**Affiliations:** Singapore Immunology Network (SIgN), Agency of Science Technology and Research (A*STAR), 8A Biomedical Drive, Singapore 138648

**Keywords:** Meiotic recombination, non-crossover recombination, genetic evolution, neutral evolution, Lamarckian evolution, transgenerational epigenetic inheritance

## Abstract

Meiotic “point recombination” refers to homologue recombination events affecting only individual SNPs. Driven mostly by gene conversion, it is common process that allows for a gradual adaptation and maturation of haplotypes during genetic evolution. In contrast to crossover recombination it is not tied to predetermined recombination sites and therefore assumed to occur largely randomly. Our analysis of archaic human haplotypes however revealed striking differences in the local point recombination rate. A linkage-study of 1.9 million SNPs defined by the sequence of *denisovan* hominids revealed low rates in introns and quiescent intergenic regions but high rates in splice sites, exons, 5’- and 3’-UTRs, and CpG islands. Correlations with ChIP-Seq tracks from ENCODE and other public sources identified a number of epigenetic modifications, that associated directly with these recombination events. A particularly tight association was observed for 5-hydroxymethylcytosine marks (5hmC). The mark was enriched in virtually all of the functional regions associated with elevated point recombination rates, including CpG islands and ‘poised’ bivalent regions. As intermediate of oxidative demethylation, 5hmC is also a marker of recently opened gene loci. The data, thus, supports a model of ‘guided’ evolution, in which point recombination is directed by 5hmC marks towards the functionally relevant regions.

## Introduction

Genetic evolution is based on the constant adaptation of the genomic sequence by mutation and genetic recombination. However, only a small fraction of the genome is directly associated with function: nearly 70% is quiescent and function-related regions such as exons, promoter and enhancer elements are condensed into less than 5% of the total genomic space (Roadmap Epigenomics Consortium et al. 2015). On the molecular level, ‘function’ is reflected in the state of the chromatin and the epigenetic patterns, namely histone- and DNA-modifications. At least in principle, these structural and epigenetic elements could therefore form a scaffold to direct the mutation and/or recombination machinery towards active genes and regions related to gene- and chromatin-function.

In order to determine if this epigenetic link actually exists, we focused on meiotic point recombination. The term refers to the allelic swap of a single SNP by homologue recombination. Contrary to point mutations, which are mostly caused by random replication errors, it requires an extensive machinery of enzymes and factors. It is typically driven by gene conversion although a certain fraction might also derive from multiple rounds of uneven crossover (CO) recombination. In contrast to CO recombination, gene conversion is not tied to predetermined breakpoints defined by PRDM9 binding sites (Baudat et al. 2010). It is also part of the regular DNA damage repair (DDR) and thus assumed to occur evenly thorough the entire genome. It may, thus, represent a very effective mechanism to facilitate a gradual adaptation of haplotypes in response to persisting environmental pressure (Sabeti et al. 2002).

For this study, we analysed the point recombination events shaping the human genome during the past 500,000+ years. Recombination rates in archaic haplotypes, defined by the sequenced genomes of *denisovan* hominids (Meyer et al. 2012a), were estimated using the genotype information of modern humans (The 1000 Genomes Project Consortium 2010; The 1000 Genomes Project Consortium 2012). The analysis revealed high recombination rates in functionally relevant regions such as exons, 5’- and 3’-UTR, open chromatin and CpG islands but low rates in introns and intergenic regions. Correlation with ChIP-seq tracks from ENCODE and other published sources indicated a clear association of the rate with open chromatin and a number of epigenetic marks. The strongest association was detected for 5-hydroxymethylcytosine (5hmC), which makes it a promising candidate to facilitate a targeted or even ‘guided’ form of genetic evolution.

## Results

### Archaic linkage blocks

The analysis was carried out with the genotype data of 99 individuals of the Luhya people from Webuye, Kenya (LWK) (The 1000 Genomes Project Consortium 2010; The 1000 Genomes Project Consortium 2012). Only those SNPs were considered, for which the derived alleles are found in the genome of the *denisovan hominid* (Meyer et al. 2012a) while their ancestral allele is still present in chimpanzees (Chimpanzee Sequencing and Analysis Consortium 2005). This resulted in a dataset of 1.9 million ‘archaic’ SNPs. As an East-African population, the LWK genomes were seemingly unaffected by late interbreeding with *denisova h*. (Meyer et al. 2012a). Most of the derived alleles of these archaic SNPs should therefore have been introduced prior to the separation of *denisova h*. and *h. sapiens*. This provides a time window of more than 500.000 years in which the alleles could be rearranged by point recombination (supplemental figure 1).

To analyse point recombination events, it was necessary to determine the sequence of the haplotype pairs that existed at the start of the recombination window. This was provided directly by the genomic sequence of *denisova* and chimpanzee, as the string of derived alleles preserved in *denisova h*. represented the evolutionary younger derived haplotype (defined here as ‘archaic haplotype’), while the corresponding ancestral alleles of the chimpanzee genome were defined as the matching ancestral haplotype (figure 1a). Due to CO recombination, they are now broken down into discrete linkage blocks. In the case of the LWK cohort, 1,897,400 archaic SNPs were arranged in 237,312 discrete blocks, each comprising 2 to more than 500 SNPs (table 1).

**FIGURE 1:**
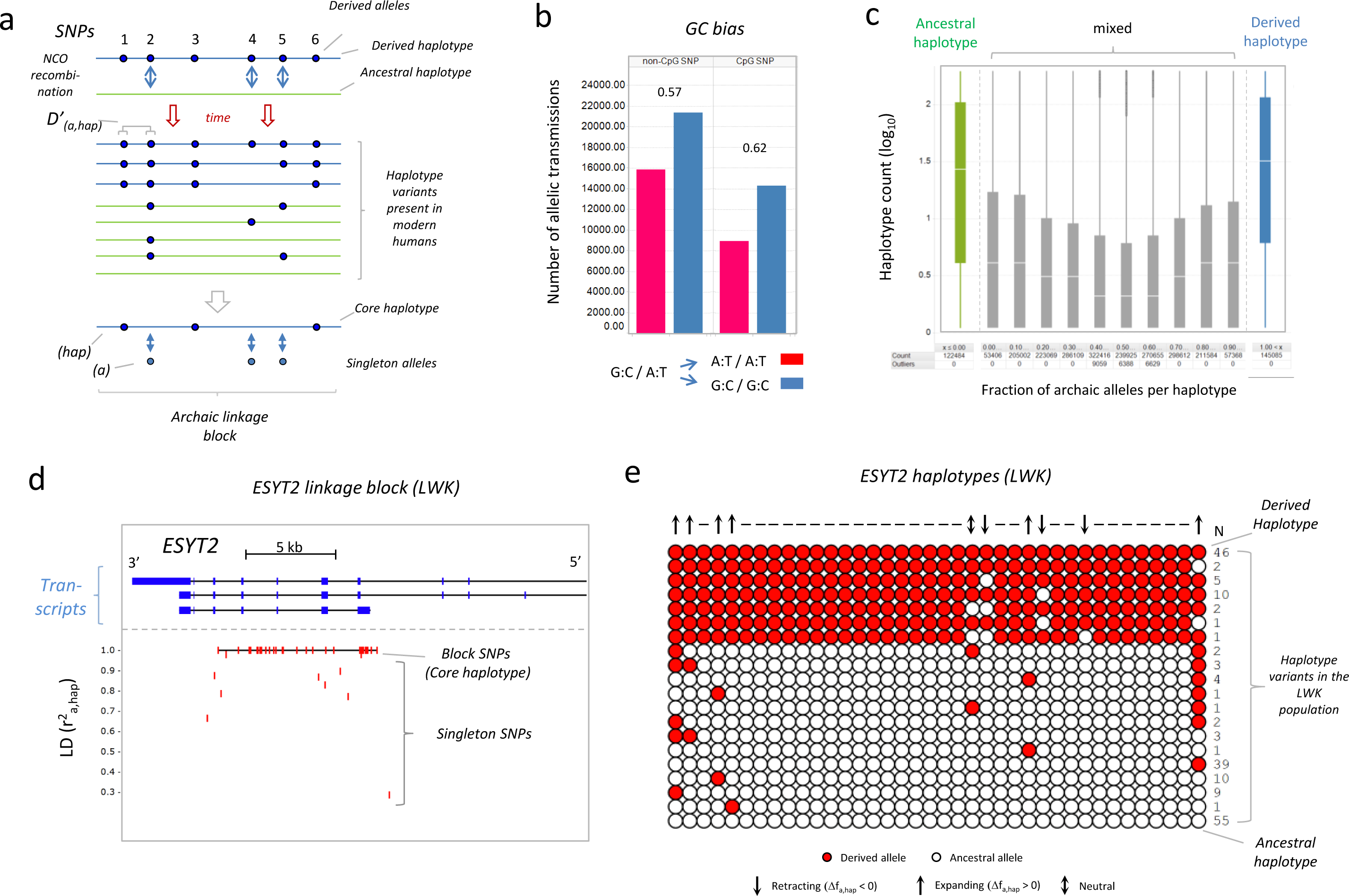
Archaic linkage blocks. **(a) Formation of archaic linkage blocks**. Archaic SNPs were formed by point mutations introduced into the genomes of early hominids (see supplemental figure 1). Genetic recombination over the past 500,000+ years between the derived haplotype (blue), consisting of the derived alleles of these SNPs (solid circles), and the corresponding ancestral haplotype (green) then created the archaic linkage blocks present in modern humans. In the given example SNPs 1, 3 and 6 are still in perfect linkage (block SNPs). The derived alleles of these SNPs form the core haplotype (hap), which represents the remaining fragment of the ‘archaic’ derived haplotype. SNPs 2, 4 and 5 are singletons affected by NCO recombination. Their association to the derived haplotype is indicated by the linkage disequilibrium (D’_a,hap_) between their derived alleles (a) and the core haplotype (hap). **(b) GC bias of point recombination**. The transmission of A:T (red) or G:C (blue) alleles is shown for a subset of archaic SNPs heterozygous for A:T / G:C alleles (see methods section). Separate plots are shown for non-CpG and CpG SNPs; numbers represent the respective GC bias. **(c) Relative frequency of archaic and ancestral haplotypes**. The block diagram illustrates the absolute number of haplotypes of archaic linkage blocks that are representing perfectly preserved ancestral (green), perfectly preserved derived (blue) or mixed haplotypes (grey). The latter consist of both derived and ancestral alleles and are binned according to the relative fraction of derived alleles per haplotype. Horizontal lines represent the median. **(d) Example of an archaic linkage block**. The panel shows the location of the SNPs of an archaic linkage block in reference to the intron/exon structure of the ESYT2 gene. As block SNPs is defined by r^2^ =1, in this visualization the association of the derived alleles to the core haplotype is indicated by the respective r^2^_a,hap_ value. The horizontal line connecting the block-SNPs indicate the location of the core haplotype; single vertical lines indicate mobility and location of the singleton SNPs. **(e) Haplotype composition of the linkage block**. The panel displays the haplotype composition of the ESYT2 linkage block provided for the 99 individuals of the LWK cohort. The presence of derived alleles (red circles) and ancestral alleles (white circles) for each haplotype variant is indicated. Numbers indicate the absolute frequency of the haplotype in the cohort; downward and upward arrows on top of the plot respectively indicate retracting and expanding derived alleles, a double arrow indicates a neutral allele that switched the position in several haplotypes without affecting its allele frequency; horizontal lines refer to block SNPs.

**TABLE 1:**
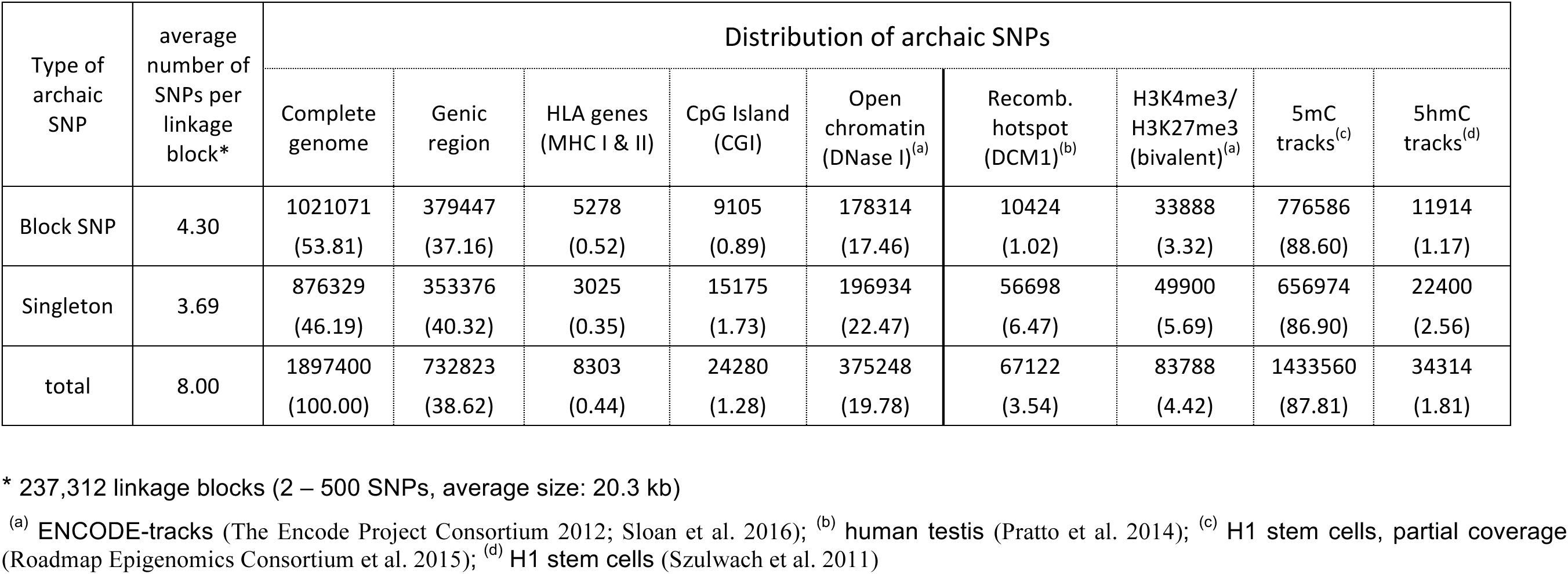
Archaic SNPs in the LWK population (1000 genome project). The table indicates the average number of block SNPs and singletons in the archaic linkage blocks of LWK and summarizes the distribution of the archaic alleles in structurally defined genomic sub-regions as well as in regions defined by ChIP-seq-tracks. Numbers indicate the absolute count of archaic SNPs detected in the region; numbers in brackets indicate the percentage in reference to the total amount of the respective SNP type.

| Type of archaic SNP | average number of SNPs per linkage block* | Distribution of archaic SNPs |  |  |  |  |  |  |  |  |
| --- | --- | --- | --- | --- | --- | --- | --- | --- | --- | --- |
|  |  | Complete genome | Genic region | HLA genes (MHC I & II) | CpG Island (CGI) | Open chromatin (DNase I)<sup>(a)</sup> | Recomb. hotspot (DCM1)<sup>(b)</sup> | H3K4me3/H3K27me3 (bivalent)<sup>(a)</sup> | 5mC tracks<sup>(c)</sup> | 5hmC tracks<sup>(d)</sup> |
| Block SNP | 4.30 | 1021071<br>(53.81) | 379447<br>(37.16) | 5278<br>(0.52) | 9105<br>(0.89) | 178314<br>(17.46) | 10424<br>(1.02) | 33888<br>(3.32) | 776586<br>(88.60) | 11914<br>(1.17) |
| Singleton | 3.69 | 876329<br>(46.19) | 353376<br>(40.32) | 3025<br>(0.35) | 15175<br>(1.73) | 196934<br>(22.47) | 56698<br>(6.47) | 49900<br>(5.69) | 656974<br>(86.90) | 22400<br>(2.56) |
| total | 8.00 | 1897400<br>(100.00) | 732823<br>(38.62) | 8303<br>(0.44) | 24280<br>(1.28) | 375248<br>(19.78) | 67122<br>(3.54) | 83788<br>(4.42) | 1433560<br>(87.81) | 34314<br>(1.81) |

\* 237,312 linkage blocks (2 – 500 SNPs, average size: 20.3 kb)
<sup>(a)</sup> ENCODE-tracks (The Encode Project Consortium 2012; Sloan et al. 2016); <sup>(b)</sup> human testis (Pratto et al. 2014); <sup>(c)</sup> H1 stem cells, partial coverage (Roadmap Epigenomics Consortium et al. 2015); <sup>(d)</sup> H1 stem cells (Szulwach et al. 2011)

While the majority of the archaic SNPs was still arranged in the original allelic order (‘block SNPs’), 876,329 SNPs were identified as singleton SNPs (affected by point recombination). In line with the GC bias reported for gene conversion (Lesecque et al. 2013; Lachance and Tishkoff 2014; Glémin et al. 2015; Williams et al. 2015), 57% of the non-CpG singleton SNPs had transmitted G:C over A:T (figure 1b). For CpG SNPs this bias increased to 62%, confirming that the allelic rearrangements in our database are indeed mostly the result of gene conversion events.

In order to quantify the recombination rates of the singleton SNPs, it was necessary to define a reference. On average, each block consisted of about 4 block SNPs and 4 singleton SNPs (table 1). The derived alleles of the block SNPs (r^2^ = 1) form the core haplotypes representing the remaining fragments of the original archaic haplotype (figure 1a). Intact versions of archaic and ancestral haplotypes are in fact the most common haplotype variants found in archaic linkage blocks (figure 1c). For each singleton SNP the point recombination rate was therefore assessed by analysing the linkage disequilibrium (LD) of their derived allele (a) in reference to the respective core haplotype (hap) (figure 1a). In the following, 1 - D’^2^_a,hap_ was therefore used as a measure for the ‘allelic mobility’. Contrary to r^2^, D’ is independent of the allele frequency and hence a more accurate proxy of the recombination rate. One example of an archaic linkage block is shown in figure 1d.

### Impact of fitness and function

In an initial attempt to identify links between point recombination rate and ‘function’ we correlated the allelic mobility of our archaic SNPs (1 - D’^2^_a,hap_) with the scores of two algorithm-based databases. The ‘Fitness consequence of functional annotation scores’ (fitCons) (Gulko et al. 2015) are indicative of the putative fitness contribution of a SNP, while the “Genome-wide annotation of variants”-scores (GWAVA) (Ritchie et al. 2014) predict the functional impact of genetic variants. Although direct correlations with the entire data set were weak (rho values of the Spearman’s rank correlation were only 0.07 and 0.01 for GWAVA- and fitCons-scores, respectively, (data not shown)), a correlation with the average scores of mobility bins revealed a significant trend for GWAVA-scores (figure 2a). The same result was also obtained when the correlation was carried out with the allele frequency of the archaic SNPs (figure 2b). Although the absolute allele frequency of derived alleles (f_a_) did not yield any association, the correlation with Δf_a,hap_, defined by the difference in frequency of derived allele and core haplotype, revealed a positive association with GWAVA-scores, evident for both expanding (Δf_a,hap_ > 0) and retracting alleles (Δf_a,hap_ < 0).

**FIGURE 2:**
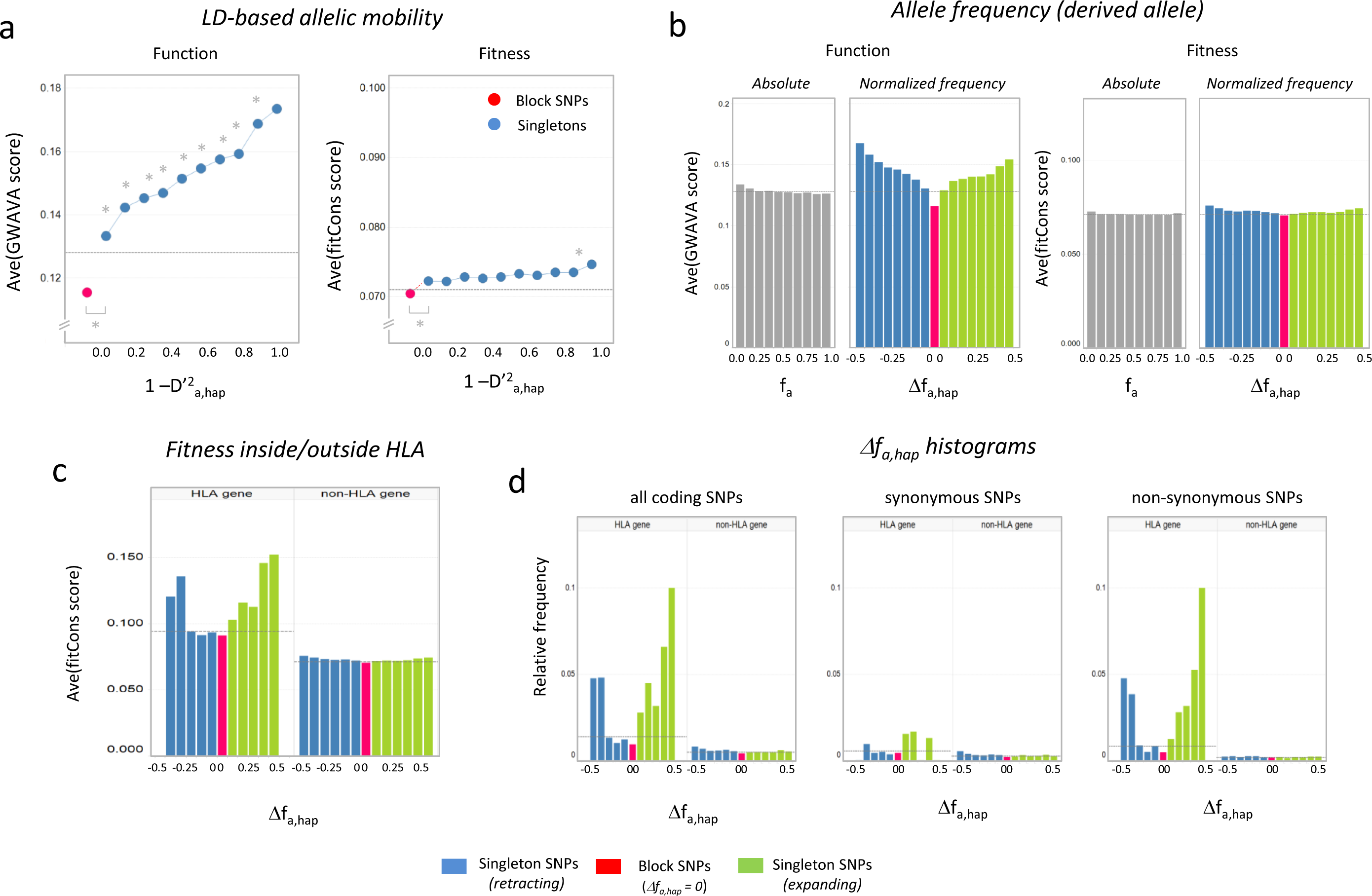
Correlation of the point recombination rate with function and fitness. **(a) Allelic mobility**. The correlation of the LD-based mobility parameter 1 - D’^2^_a,hap_ with function-associated GWAVA scores (left panel) and fitness-associated fitCons scores is shown. The plot displays the average scores for 1.9 million archaic SNPs divided into 1.0 million block SNPs (red dot) and about 900,000 singleton SNPs (blue dots). The singleton SNPs were binned according to the 1 - D’^2^_a,hap_ value defined for the LWK population. Dashed horizontal lines indicate the average scores of the entire set of archaic SNPs. Asterisks indicate significant differences between data point pairs (p < 0.05; Mann-Whitney U test). **(b) Allele frequency**. Allele frequencies of the archaic SNP set are plotted against the scores of the GWAVA- (left panels) and the fitCons-database (right panel). Separate plots are shown for the absolute frequency f_a_ of derived alleles (grey bars) and the normalized frequency Δf_a,hap_, representing the difference in frequency between allele and core haplotype. Blue bars represent retracting derived alleles (Δf_a,hap_ < 0), green bars expanding derived alleles (Δf_a,hap_ > 0); red bars comprise neutral alleles (Δf_a,hap_ = 0), mostly representing the alleles of the core haplotypes. Each bar indicates the average score of the respective allele frequency bin; dashed lines indicate the average score of the entire dataset. **(c) Fitness inside and outside the HLA gene region**. The correlation of fitCons scores with the normalized allele frequency Δf_a,hap_ is shown for derived alleles located within or outside of HLA genes (MHC class I and class II). **(d) Δf_a,hap_ histograms of coding alleles**. The histograms display the relative frequency of coding SNPs binned according to their normalized allele frequency Δf_a,hap_. A comparison is shown for derived alleles located within or outside of HLA genes, displaying the relative frequencies for all coding SNPs (left panel), synonymous coding SNPs (middle panel) and non-synonymous coding SNPs (right panel).

In contrast to GWAVA-scores, fitCons scores showed no association with neither 1 – D’^2^_a,hap_ (figure 2a) or Δf_a,hap_ (figure 2b). However, a strong association with Δf_a,hap_ was detected, when the allele set was restricted to SNPs located in the MHC genes of the HLA region (figure 2c). Histograms displaying the relative frequency of coding SNPs in Δf_a,hap_ bins confirmed that the allelic expansion is indeed driven by selection: the correlation was evident for non-synonymous but not for synonymous SNPs (Figure 2d). Outside of the HLA region only marginal effects were detected and no bias towards non-synonymous coding SNPs was evident. This result is thus in line with the prevailing model of neutral evolution, allowing fitness-driven selection essentially only for the alleles of MHC genes (Klein 1996).

### Chromatin and genic sub-regions

To further characterize the link between point recombination and ‘function’, we correlated the mobility parameter 1 - D’^2^_a,hap_ with gene regions and other annotated information related to the genome structure (table 1, figure 3). In a correlation with the distance to the closest gene, particularly sharp peak of mobility was detected proximal to the 5’ boundary and, to a weaker extent, also at the 3’ boundary of annotated gene regions (figure 3a). Further delineation of genic sub-regions corroborated this finding with high mobility values observed in the 5’- and 3’-UTR as well as in the respective flanking regions located upstream and downstream of the genes (figure 3b). Within the gene sub-regions, allelic motilities were particularly high in exons, followed by splice sites and introns (figure 3b). In intergenic regions, the mobility dropped with increasing distance to the gene boundaries (figure 3b). Correlation with ENCODE tracks of DNase I sensitivity (The Encode Project Consortium 2012; Sloan et al. 2016) further revealed an increase in allelic mobility in regions of open chromatin (Figure 3 c). This applies particularly for alleles in longer open stretches, indicated by a sharp mobility increase for alleles located more than 500bp from the boundaries (comprising less than 0.5% of the SNPs).

**FIGURE 3:**
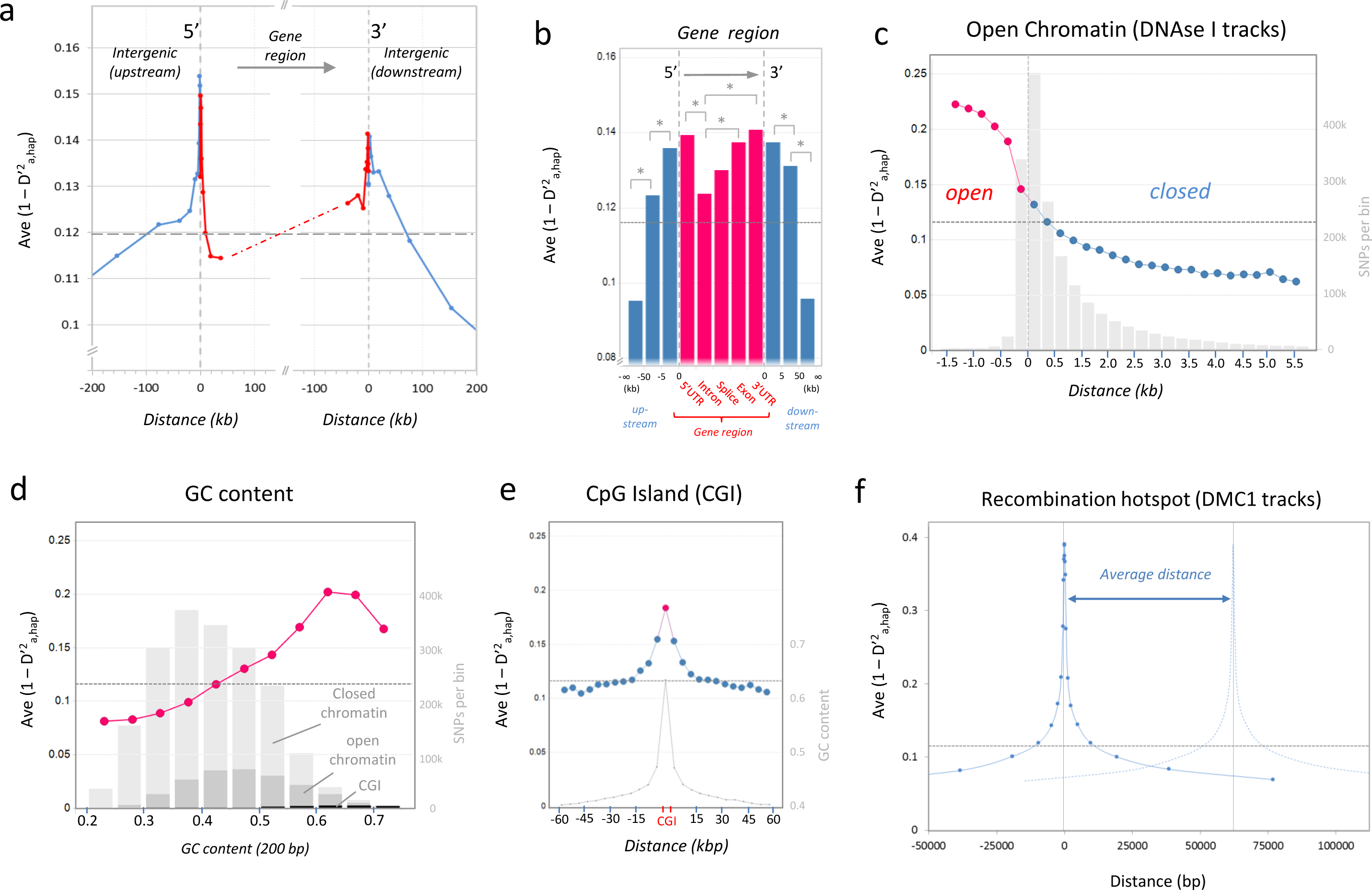
Correlation of the point recombination rate with structural genomic parameters. **(a) Gene boundaries**. The point recombination rate of 1.9 million archaic SNPs, expressed as average allelic mobility (1 - D’^2^_a,hap_), is plotted in reference their distance to the boundary of the closest annotated gene. Peaks in mobility are evident at both 5’- and 3’- boundaries (indicated by dashed vertical lines). Gene regions are marked in red, intergenic regions in blue; dashed horizontal line represents the average mobility of the entire set of archaic alleles. **(b) Genic sub-regions**. Gene regions (red) were further delineated into their sub-regions (5’UTRs, introns, splice sites, exons and 3’UTRs). Intergenic regions (blue) were divided according to their distance to the boundaries both upstream and downstream of the genic region (0-5 kb, 5-50 kb, >50kb). Bars represent the average allelic mobility (1 - D’^2^_a,hap_) of archaic alleles in the respective region. Asterisks indicate significant differences between the indicated bars (p < 0.05; Mann-Whitney U test). **(c) Open chromatin**. The line chart depicts the association of the point recombination rate with open chromatin. The average allelic mobility (1 - D’^2^_a,hap_) is plotted in reference to the distance of the alleles to the boundary of open chromatin (defined by ENCODE tracks of DNase I sensitivity). Grey bars indicate the number of archaic SNPs in the respective distance-bin. **(d) Local GC content**. The line chart indicates the association of the point recombination rate with the local GC content. The average allelic mobility (1 - D’_a,hap_) is correlated with the relative GC content of a 200bp window surrounding the SNP. Grey bars indicate the number of archaic SNPs in the respective GC content-bin subdivided into regions of closed chromatin (light grey), open chromatin (middle grey) or CpG islands (CGI; dark grey). **(e) CpG islands**. The coloured line chart indicates the NCO recombination rate in CpG islands (CGI). The average allelic mobility (1 - D’^2^_a,hap_) is plotted in reference of their distance to the boundary of CGI as generated by UCSC Genome Browser. The plot covers alleles located inside (red) and outside (blue) of CGI; grey peak represents the GC content for the respective distance bin. **(f) Meiotic recombination hotspots**. The solid line represents the average allelic mobility (1 - D’^2^_a,hap_) plotted in reference to the distance of the alleles to meiotic recombination hotspots (defined by DMC1 ChIP-Seq tracks of human testis (Pratto et al. 2014)). The dashed line of the second peak represents the hypothetical mobility in reference to a second recombination hotspot located at the average distance of about 67 kb.

It is already well established that the non-crossover (NCO) recombination rate increases with the local GC content (Halldorsson et al. 2016). In line with these reports also the allelic mobility of our archaic SNPs was found to be directly associated with this parameter (Figure 3d). The curve, however, does not reflect a simple linear correlation but rather reaches a maximum at a GC content of about 0.6-0.65. This GC content characteristic for CpG islands (CGI) and a steep rise in CGI regions was in fact observed when plotting the mobility parameter in reference to the distance to the closest CGI (Figure 3e).

A strong influence on the mobility was also excepted to be observed for meiotic recombination hotspots. While these hotspots primarily represent pre-determined breakpoints for CO recombination, they often also serve as initiation sites of NCO recombination (Jeffreys and May 2004; Odenthal-Hesse et al. 2014; Williams et al. 2015). DMC1 is a homolog of the bacterial strand exchange protein RecA and acts as a specific marker of these hotspots (Bishop et al. 1992; Bishop 1994). The mobility correlation with ChIP-Seq tracks of human testis (Pratto et al. 2014) in fact revealed extremely strong 1 - D’^2^_a,hap_ values for alleles located proximal to the binding sites of DMC1 (Figure 3f). The effect, however, was very local: at a distance of approximately 1,000 bp, the mobility declined already to half. With an average spacing of about 65kb, the influence of hotspots is, therefore, restricted to less than 5% of the SNPs (table 1).

### Influence of epigenetic histone marks

To determine if the point recombination rate is influenced by epigenetic histone modifications, we used the ChIP-Seq tracks of provided by ENCODE (H3K27me3, H3K4me1, H3K4me3, H3K4me2, H3K9ac, H3K27ac, H3K79me2, H3K36me3, H3K9me3). In order to assess their impact independent of the influence of the genomic location, separate mobility correlations were carried out for seven sub-regions (intergenic, < 5kb upstream, 5’ UTR, intronic, exonic, 3’-UTR, and > 5kb downstream regions).

Notably, all of the analyzed histone marks showed some form of association with the allelic mobility (figure 4a). Depending on the general direction, the impact of the modification could be classified either as ‘recombination-promoting’ (H3K27me3 ≫ H3K4me1, H3K9ac, > H3K4me3, H3K4me2 > H3K27ac) or ‘recombination-repressing’ (H3K9me ≫ H3K36me3, H3K79me2). The strongest positive association was observed for H3K27me3, while the strongest negative association was observed for H3K9me3.

**FIGURE 4:**
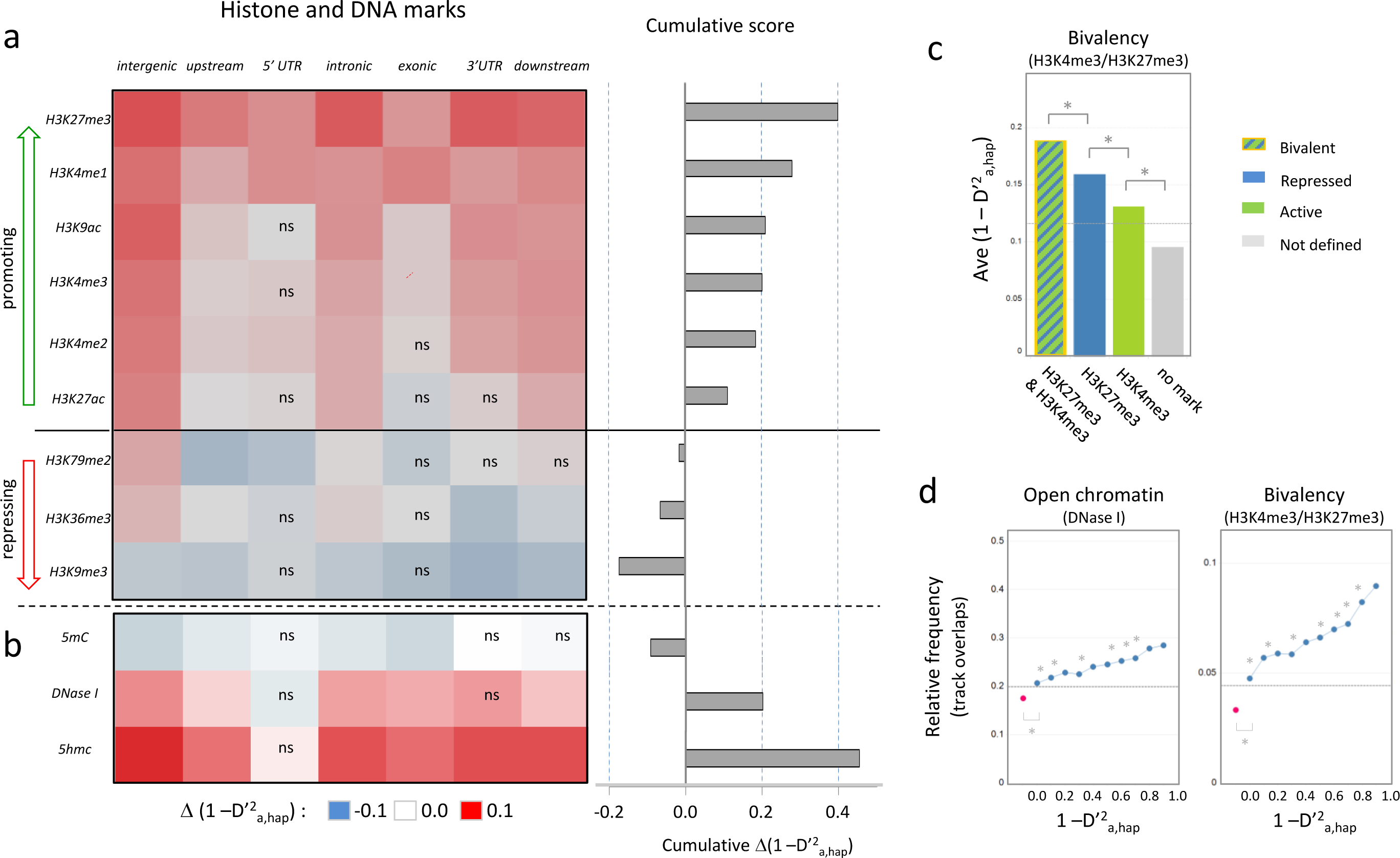
Correlation of the point recombination rate with epigenetic marks. **(a) Histone marks**. The impact of epigenetic marks on the point recombination rate was assessed by determining the average allelic mobility (1 - D’^2^_a,hap_) for alleles located on tracks of histone marks (H3K27me3, H3K4me1, H3K4me3, H3K4me2, H3K9ac, H3K27ac, H3K79me2, H3K36me3, H3K9me3). The data set was further divided based on the location of archaic alleles in genic sub-regions (intergenic, upstream > 5 kb, 5’ UTR, intronic, exonic, 3’ UTR, downstream < 5 kb). Each combination of histone marks and genic sub-regions were tested using MannWhitney U test to determine if the overlap with the histone mark led to a significant difference in the allelic mobility. Left panel: The colour code in the heat map indicates the deviation from the average the allelic mobility Δ(1 - D’_a,hap_) of the respective sub-region, ranging from −0.1 (blue) to 0.1 (red); ‘ns’ indicates a non-significant association; solid horizontal line separates recombination promoting marks (green arrow) from recombination repressing marks (red arrow). Right panel: The bars represent the cumulative mobility score for each of the marks over all sub-regions. The score was calculated by adding up all significant Δ(1 - D’^2^_a,hap_) of the genic subregions. ChIP-Seq tracks of histone marks were compiled from ENCODE data of various tissues and cell lines. **(b) DNase I and epigenetic DNA marks**. The correlation was carried out as in (a) except that allelic mobility was correlated with the two DNA marks, 5-methylcytosine (5mC) and 5-hydroxymethylcytosine (5hmC), as well as for open chromatin defined by DNase I sensitivity. All tracks were derived from the embryonic stem cell H1 (Szulwach et al. 2011; Roadmap Epigenomics Consortium et al. 2015). **(c) Bivalent regions**. The average allelic mobility (1 - D’^2^_a,hap_) is shown for alleles located in bivalent regions characterized by overlapping tracks of both H3K4me3 and H3K27me3 (hatched bar), univalent regions characterized by overlapping tracks of either H3K27me3 (blue bar) or H3K4me3 (green bar) or regions without overlap to the tracks of any of the two marks (grey bar). Asterisks indicate significant differences between the indicated bars (p < 0.05; Mann-Whitney U test). **(d) Mobility correlation with open chromatin and bivalent state**. The association of the allelic mobility (1 - D’^2^_a,hap_) with open chromatin (left panel) and bivalent marks (right panel) is shown. The dots indicate the relative frequency of 1.9 million archaic alleles divided into 1 million block SNPs (red dot) and about 900,000 singleton SNPs (blue dots) located on published tracks of the respective mark. The singleton SNPs were binned according to their mobility; the tracks were obtained from ENCODE; dashed line represents the average frequencies of the entire set of archaic alleles. Asterisks indicate significant increase in overlaps with increased mobility between data point pairs (p < 0.05; Chi-square test).

While the histone marks H3K27me3, H3K4me1 and H3K9ac showed strong positive associations in various different sub-regions, particularly high mobility values were detected in bivalent regions tagged by both H3K27me3 and H3K4me3 marks (figure 4c). Compared to these, the recombination rate in regions marked only by H3K27me3 was significantly lower, and regions tagged solely by H3K4me3, exhibit only a minor increase compared to the genome-wide average. H3K4me3 is associated with active genes, while H3K27me3 is indicative of transcriptionally repressed chromatin. Bivalent histone-modifications are therefore hypothesized to 'poise' silenced genes for rapid activation (Harikumar and Meshorer 2015), a state, which according to this data, seems to prime also meiotic point recombination.

### Influence of epigenetic DNA marks

The observation that bivalency has a stronger impact on the mobility than DNase I-sensitivity suggests that presence of epigenetic marks, rather than just the physical accessibility, bear a controlling influence on the recombination rate (figure 4d, supplemental figure 2a). However, as variety of different histone modifications also showed similar correlations, it appeared more likely that, instead of individual histone marks, the resulting structural state of the chromatin or associated DNA-marks might actually be responsible for the effects. 5-methyl-cytosine (5mC) is known to be linked with closed state of chromatin, while 5-hydroxymethyl-cytosin (5hmC), as intermediate of oxidative C-demethylation, is indicative of recently opened chromatin. As both tracks were not available from the ENCODE database, they were downloaded together with matching DNase I-sensitivity tracks for the embryonic stem cell H1 from other sources (Szulwach et al. 2011; Roadmap Epigenomics Consortium et al. 2015).

Similar to the histone marks, also DNAse I and the two DNA marks showed very clear associations to the mobility (figure 4b). This applied for the DNase I sensitivity, and in an inverse relation, also by the 5mC mark. Most pronounced, however, was the correlation with 5hmC. In virtually all analyzed sub-regions the gain in mobility was the highest when the alleles overlapped with 5hmC tracks. The cumulative score of 5hmC in fact eclipsed the scores of all other tested marks, suggesting 5hmC to be the best candidate for controlling the meiotic point recombination rate (figure 4b). The direct comparison between 5mC and 5hmC in fact revealed a more than 4fold increase in the mobility-dependent fold change compared to 5mC (figure 5a, supplemental figure 2b).

**FIGURE 5.**
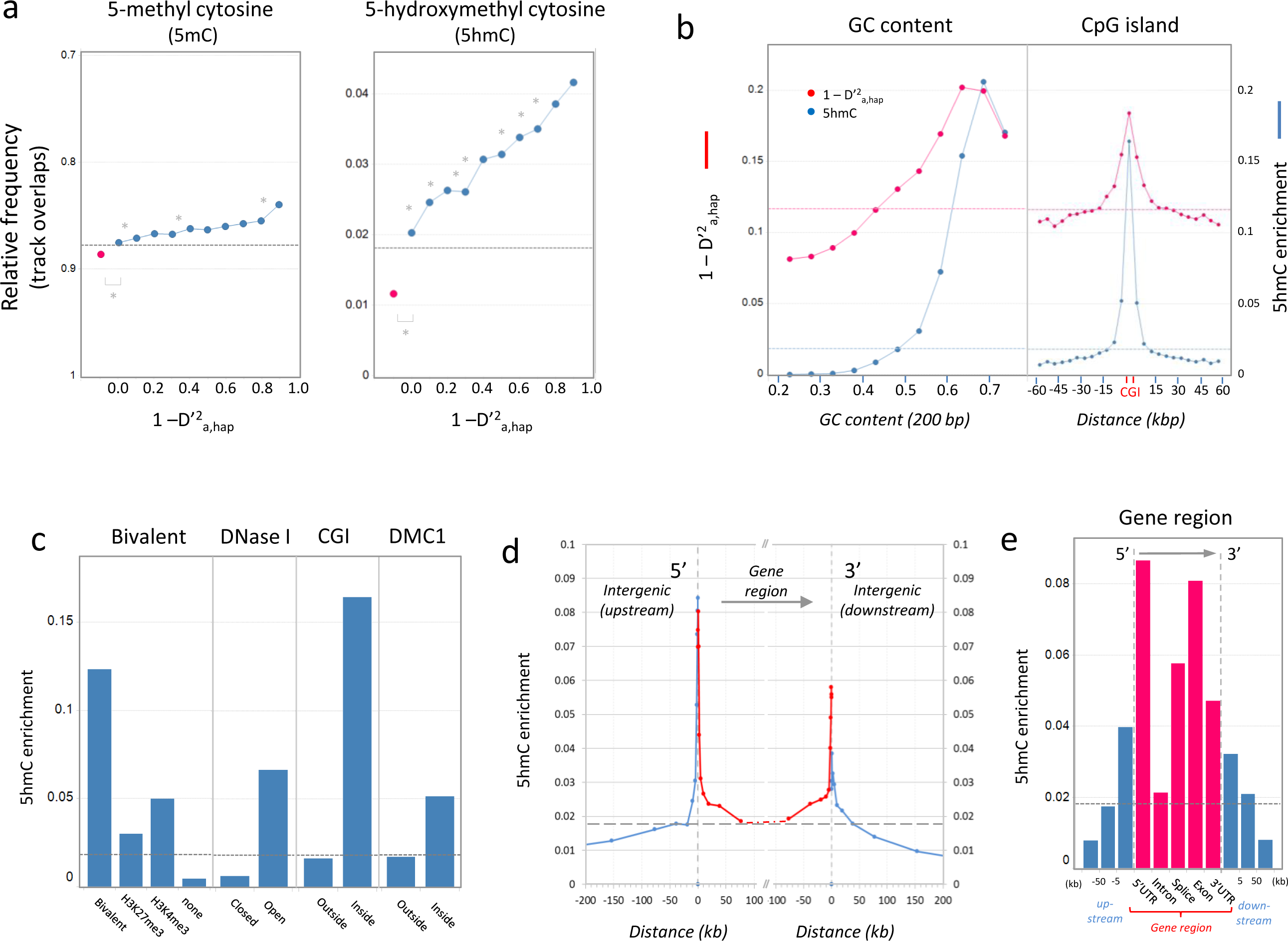
Local enrichment and rate-association of 5hmC marks. **(a) Mobility correlation with 5mC and 5hmC**. The association of the allelic mobility (1 - D’_a,hap_) with 5mC (left panel) and 5hmC (right panel) is shown. The dots indicate the relative frequency of 1.9 million archaic alleles divided into 1 million block SNPs (red dot) and about 900,000 singleton SNPs (blue dots) located on published tracks of the respective mark. The singleton SNPs were binned according to their mobility; published tracks were obtained for the stem cell line H1 (Szulwach et al. 2011; Roadmap Epigenomics Consortium et al. 2015); dashed line represents the average frequencies of the entire set of archaic alleles. Asterisks indicate significant increase in overlaps with increased mobility between data point pairs (p < 0.05; Chi-square test).**(b) Recombination rate and 5hmC enrichment of in GC-rich regions**. The plots display the NCO recombination rate expressed as 1 - D’^2^_a,hap_ (red lines) and the relative enrichment of 5hmC marks (blue lines) in reference to the local GC-content (left panel) or the distance to CpG islands (right panel). The mobility of 1.9 million archaic SNPs was calculated for LWK; the 5hmC enrichment of the bin was determined by calculating the relative frequency of archaic alleles overlapping with published 5hmC tracks of H1 cells (Szulwach et al. 2011); dashed lines represent the average values for the entire data set. **(c) Enrichment of 5hmC in genomic sub-regions**. The local 5hmC enrichment is shown for bivalent regions (“bivalent”), open chromatin (“DNase I”), CpG islands (“CGI”) and meiotic recombination hotspots (“DMC1”). The location of bivalent regions (H3K27me3 & H3K4me3) and open chromatin (DNase I) was defined by ENCODE tracks; the location of CGI was defined by UCSC Genome Browser. Hotspot association was defined by the overlap with DMC1 tracks from human testis (Pratto et al. 2014). For each bar combination, the difference is significant (p < 0.05; Chi-square test). **(d) Gene boundaries**. The local 5hmC enrichment is plotted in reference to the distance to the boundary of the closest annotated gene. Peaks are evident at 5’ or 3’ boundaries (indicated by dashed vertical lines). Genic regions are marked in red, intergenic regions in blue; dashed horizontal line represents the average overlap with 5hmC tracks of the entire set of archaic alleles. **(e) Genic sub-regions**. The local 5hmC enrichment is shown for genic sub-regions (red: 5’ UTR, intron, splice site, exon and 3’ UTR) and intergenic regions (blue) binned according to the distance to the respective gene boundary (0-5kb, 5-50kb, <50kb). Horizontal dashed line represents the average 5hmC-overlap of the entire set of archaic alleles. For each bar combination, the difference is significant (p < 0.05; Chi-square test).

The close link between 5hmC and point recombination was evident in virtually all of the parameters shown in this study to influence the rate. Plots representing the association with the GC content revealed similar maxima curves for both 5hmc enrichment and mobility (figure 5b, left panel). The peak in the 5hmC amounts in CpG islands closely reflected the sharp increase observed also for the mobility (figure 5b, right panel), and, in line with the increase in mobility, 5hmC was also strongly enriched in bivalent regions, open chromatin, CpG islands (CGI), and DMC1 tracks (figure 5c). The 5hmC density spiked at the 5’- and, to a lesser extent, the 3’-boundaries of genes (figure 5d), closely mirroring the mobility profile shown figure 3a. Also in genic sub-regions, the 5hmC density was highest in 5’-UTR with gradually decreasing values in intergenic regions and was low in introns, increased in splice sites and was highest in exons (figure 5e; compare figure 3b). Thus, 5hmC seems to be prominently present in all the functionally relevant loci that, according to this study, are also associated with an increased point recombination rate.

### Mobility and 5hmC content in epigenetic sub-regions

To further evaluate the impact of the local epigenetic environment on the point recombination rate we used an independent data set that allows to divide the archaic SNP set into epigenetically defined regions. The NIH Roadmap Epigenomics Consortium (Roadmap Epigenomics Consortium et al. 2015) provides comprehensive data on the epigenomes of 111 tissues, cells and cell lines. Based on the distribution of histone marks and the state of the chromatin, each of these reference epigenomes is divided into 15 sub-regions (supplemental figure 3). These regions represent active promoters (TssA, TSSAFlnk), transcribed regions (TxFlnk, Tx, TxWk), active enhancers (EnhG, Enh), ZNF/Repeat and heterochromatin regions (ZNFRpts, Het), bivalently ‘poised’ regulatory regions (TssBiv, BivFlnk, EnhBiv), polycomb repressed regions (RepPC, RepPCWk) and quiescent DNA (Quies). Based on this annotation the point recombination rates (as defined by allelic mobility), as well as the fraction of open chromatin (defined by DNase I and 5mC) and the overlap with 5hmC tracks could be determined independently for each of the 15 sub-regions (figure 6). Since the recombination rate is strongly compounded by recombination hotspots (compare figure 3f), the analysis was carried out separately for SNPs located outside or inside known hotspot regions (defined by DMC1 ChIP-Seq tracks of human testis (Pratto et al. 2014)).

**FIGURE 6:**
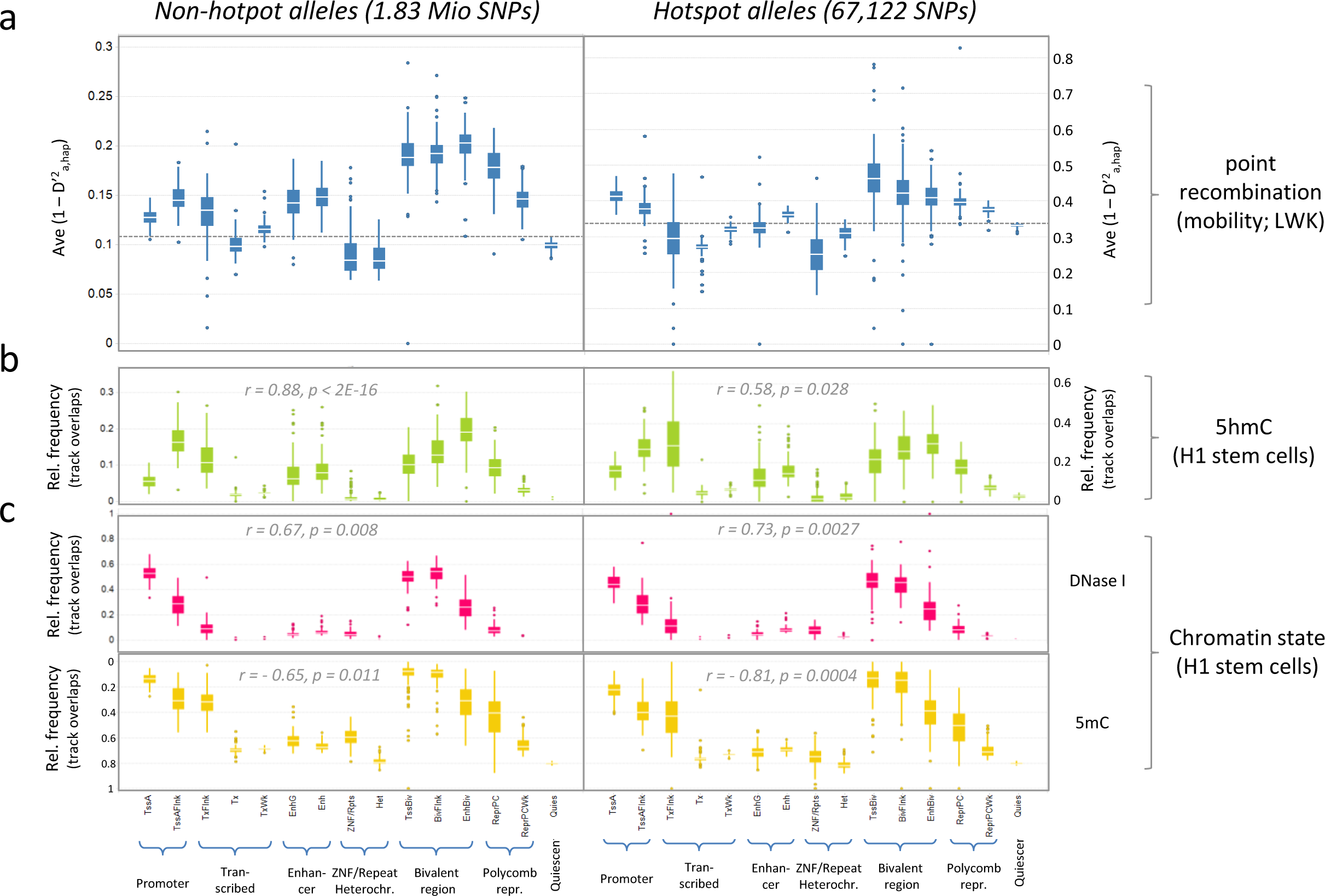
Point recombination rates in epigenetic sub-regions. NCO recombination rate, chromatin state and DNA marks are shown for 15 epigenetic sub-regions defined by the NIH Roadmap Epigenomics Consortium (Roadmap Epigenomics Consortium et al. 2015) (histone marks are shown in supplemental figure 3). TssA: active transcription start site; TssAFlnk: TssA flanking region; Tx: transcribed region; TxWk: weak transcribed region; TxFlnk: Tx flanking region; EnhG: gene-associated enhancer; Enh: enhancer; ZNF: zinc finger nuclear factor; Rpts: repeats; Het: heterochromatin; TssBiv: bivalent transcriptional start site; BivFlnk: bivalent flanking regions; EnhBiv: bivalent enhancer; ReprPC: polycomb repressed region; ReprPCWk: weak polycomb repressed region; Quies: quiescent. The calculations were carried out separately for each of 111 reference genomes using the entire set of 1.9 million archaic SNPs. Separate plots are shown for non-hotspot alleles (left panels) and alleles located at meiotic recombination hotspots (right panels). The respective assignment is based on their overlap with published DMC1 ChIP-Seq tracks of human testis (Pratto et al. 2014). **(a) Point recombination rate**. The average point recombination rate, expressed as average mobility (1 - D’^2^_a,hap_), was calculated using the mobility parameters defined for the LWK cohort. The calculation was carried out independently for each reference genome by using the respective region-annotations. The boxplots represent the variation of average mobility within these genomes; bars indicate the average of each sub-region. **(b) 5hmC marks**. The relative frequency of SNPs is shown that are located on 5hmC tracks. The tracks were defined for the H1 stem cell line (Szulwach et al. 2011). The r- and p-values refer to a Spearman rank correlation carried out between these relative frequencies and the mobility data shown in (a). **(c) Chromatin state**. The state of the chromatin was defined by DNase I tracks and 5mC marks of embryonic stem cell line H1 (Roadmap Epigenomics Consortium et al. 2015). The calculation was carried out as described in (b).

While the average allelic mobility 1 - D’^2^_a,hap_ was nearly three times higher at recombination hotspots (indicated by different y-axes in figure 6a), very similar patterns of mobility variations were evident in both sets for the epigenetically defined sub-regions (Figure 6a). The highest mobility was detected in bivalent regions representing ‘poised’ versions of promoter (TssBiv), enhancer (EnhBiv) and flanking regions (BivFlank) (supplemental figure 3). In comparison, the mobility in the corresponding active counterparts TssA, EnhG, Enh and TssAFlnk were substantially lower. Slightly lower mobility was detected in Polycomb-repressed regions, which are associated with recombination-promoting H3K27me3 marks, while the lowest mobility was detected in ZNF/Repeat and Heterochromatin regions, two regions associated with the recombination-repressing H3K9me3 mark (compare supplemental figure 4). Rather low mobility was also determined for alleles in transcribed regions (Tx, TxWk, TxFlnk), which is presumably due to the fact that they largely comprise of introns associated with low mobility.

Strikingly similar patterns emerged when comparing mobility in the various epigenetic regions (figure 6a) with the relative track-overlaps of 5hmc (figure 6b) and DNaseI, 5hmC (figure 6c). For DNase I-defined open chromatin (figure 6c) the Spearman correlation with mobility revealed r = 0.73 and p = 0.0027 for hotpot alleles and r = 67, p = 0.008 for non-hotspot alleles. For 5mC tracks the correlation was inversely mirrored but equally strong (non-hotspot alleles: r = - 0.65, p = 0.011; hotspot alleles: r = - 0.81, p = 0.0004). However, an even better match was evident for 5hmC (figure 6b). Firstly, in contrast to DNase I and 5mC, the hotspot-associated increase in mobility was directly reflected in the track overlaps with 5hmC. The frequency was increased more than 2 –fold compared to the non-hotspot regions (indicated by different y axes in figure 6b). Secondly, the calculated association between 5hmC overlap and mobility by the Spearman correlation was exceptionally strong. This applied especially for non-hotspot alleles (r = 0.88 and p = 2E-16), comprising more than 95% of the alleles. Within the hotspots, the correlation was weaker (r = 0.58 and p = 0.028) suggesting a less crucial role of the mark at CO-recombination sites. 5hmC may thus be a prime candidate to guide the ‘natural’ meiotic gene conversion process, taking place independent of predetermined recombination hotspots.

## Discussion

By analysing point recombination events in archaic human haplotypes, we established that the local recombination rates are to some extent aligned with the putative functional value of the respective genomic region. Elevated rates were observed in UTR, splice sites and exons, while in intergenic regions the rates decrease with increasing distance to the gene boundaries. Increased rates were also found in open chromatin, regions with high GC-contents and CpG islands and especially in bivalently ‘poised’ regulatory regions. While various histone marks associated with the rate in either a positive or negative way, the closest correlation was observed for 5hmC. The DNA mark was enriched in all of the functionally relevant regions associated also with increased point recombination rates.

Throughout the study D’ was used as proxy for the recombination rate. Although the parameter is mostly reflective on the number of haplotype variations present in a population, some influence of natural selection cannot be excluded. Correlations with fitness-related fitCons scores, however, revealed clear links to allelic selection only for SNPs in the HLA gene region. Also, a biased allelic expansion towards non-synonymous coding SNPs (indicative of natural selection) was evident only inside, but not outside, of this region, confirming the validity of the use of D’ as rate surrogate. As the MHC genes are the well-known exception of neutral evolution (Klein 1996), our analyses are also consistent with the concept of neutral evolution. According to this model, haplotypes are formed primarily by chance rather than be selection. As an important extension however our data suggests that the recombination bias towards functional regions may increase the probability of actually forming selectable haplotypes to drive evolution.

Variations in the local recombination rate had been reported before. Some of them are gender-specific, a phenomenon known as heterochiasmy (Mank 2009). For instance, it is well established that the recombination rate in telomeric regions is elevated in males compared to females (de Massy 2013). More importantly, a comprehensive study of the deCODE consortium of 15,257 Icelandic parent-offspring pairs indicated that, contrary to our data, human recombination appeared to be repressed within gene regions (Kong et al. 2010). The analysis, however, covered mostly alleles that had been swapped by chromosomal crossover (CO). The predetermined sites used for the double strand breaks (DSB) for are therefore primarily defined by the binding motif of the zinc finger domains of PRDM9 (Myers et al. 2008; Baudat et al. 2010; Myers et al. 2010). Recombination hotspots are enriched in bivalent regions (Zeng and Yi 2014) and the process of CO recombination itself is tightly linked to H3K4me3 marks produced by PRDM9, which acts as germ line-specific methyltransferase (Baudat et al. 2010; Berg et al. 2010; Myers et al. 2010; Parvanov et al. 2010). This seems to create specific epigenetic pattern near the PRDM1 binding site that directs especially CO recombination away from functional genomic elements (Brick et al. 2012).

In dogs, the orthologue of PRDM9 is non-functional (Munoz-Fuentes et al. 2011). The recombination studies carried out in this species revealed a pattern much more consistent with our data: elevated rates were detected in promotor regions close to the transcriptional start site and around CpG islands (Auton et al. 2013). Likewise, also in mice lacking functional PRDM9 the DSB-hotspots are enriched in promoter- and CpG-rich-regions (Brick et al. 2012). By restricting our analysis to point recombination events, we eliminated the bulk of PRDM1-based recombination events. Similar to the recombination patterns in PRDM9-deficient animals, the allelic rearrangement of isolated SNPs now targets preferably functional elements. This process, presumably driven mostly by gene conversion, is apparently guided by epigenetic marks. Point recombination strongly associated with H3K27me3 but also a number of other H3 modifications, such as H3K4me1 and H3K9ac, correlated positively with the rate and negative associations were detected for H3K36me3 and H3K9me3.

As for CO recombination (Zeng and Yi 2014), a particularly strong correlation was observed for bivalent regions marked by both H3K4me3 and H3K27me3. This was confirmed in the analysis of epigenetic sub-regions, revealing increased point recombination rates in the ‘poised’ versions of enhancer, promoter and flanking regions compared to their active counterparts. Notably, a recent study indicated that in germ-line cells, poised genes often act as ancient regulators (Choate and Danko 2016). As the lineage-specific poising correlates here with evolutionary innovations, it is intriguing that both point recombination and CO recombination, seems to be directed to bivalent regions.

In contrast to histone marks that associated only in a composite manner, open chromatin and, even more profoundly, 5hmC correlated directly with the point recombination rate. 5hmC is a unique epigenetic mark (Munzel et al. 2010) influencing chromatin structure and DNA Binding Protein (DBP) interactions (Valinluck et al. 2004; Xu et al. 2011a). It is strongly associated with bivalent chromatin (Pastor et al. 2011) as well as with promoters bearing the polycomb repressive mark, H3K27me3 (Williams et al. 2011; Wu et al. 2011a; Wu et al. 2011b; Wu and Zhang 2011b; Wu and Zhang 2011a). 5hmC is also enriched in genic regions (Stroud et al. 2011; Xu et al. 2011b) with increased levels in exons (Szulwach et al. 2011; Xu et al. 2011b), transcription factor binding sites (Stroud et al. 2011; Szulwach et al. 2011) and 5’UTRs and TSSs of genes (Pastor et al. 2011). Based on our study, the local density of the mark closely corresponds to the frequency of point recombination events: both parameters are increased in bivalent regions, open chromatin, CpG islands and recombination hotspots, they peak in exons and UTRs and follow a very similar trend in epigenetic sub-regions (bivalent ≫ polycomb-repressed ≫ active enhancer > promoter ≫ transcribed regions > quiescent > ZNF/repeats, heterochromatin).

One plausible mechanism linking the 5hmC mark to point recombination could be that it acts as a tag for induced double strand breaks. A candidate to execute this cut is ENDOG, a ubiquitous eukaryotic restriction enzyme that reportedly creates substrates for genetic recombination by cleaving 5hmC-modified DNA (Robertson et al. 2014). Studies with ENDOG -/- mice have established already a role in the somatic recombination required for immunoglobulin class switching (Zan et al. 2011). While no reports have surfaced yet that implicate 5hmC directly with meiotic recombination, in somatic cells it co-localizes with 53BP1 and *γ*H2AX, two key-components of the DNA damage repair (DDR) complex (Kafer et al. 2016). As the DDR pathway is also active in the germ line, 5hmC-directed gene conversion may therefore act as a recombination pathway parallel to the canonical CO recombination pathway, targeting only the predetermined recombination sites.

As mentioned above, the enrichment of 5hmC in exons, splice sites, UTR, enhancer or CpG islands should facilitate a ‘targeted’ evolution, in which point recombination is directed towards these function-related regions (figure 7a). However, besides acting as a static epigenetic marker, 5hmC is also a dynamic marker of recently activated loci. As a product of the cytosine-demethylation pathway (Tahiliani et al. 2009), it appears in chromatin that has newly undergone a conformational conversion from closed to open state (Pandiyan et al. 2013). Oxidative demethylation can occur in response to environmental stress, as observed, for instance, in dendritic cells exposed to *M. tuberculosis* infection (Pandiyan et al. 2013; Pacis et al. 2015). Provided, these signals are transmitted into the germ line cells, the epigenetic response could direct the recombination machinery towards the respective stress-response genes, facilitating a ‘guided’ evolution in the Lamarckian sense (figure 7b).

**FIGURE 7:**
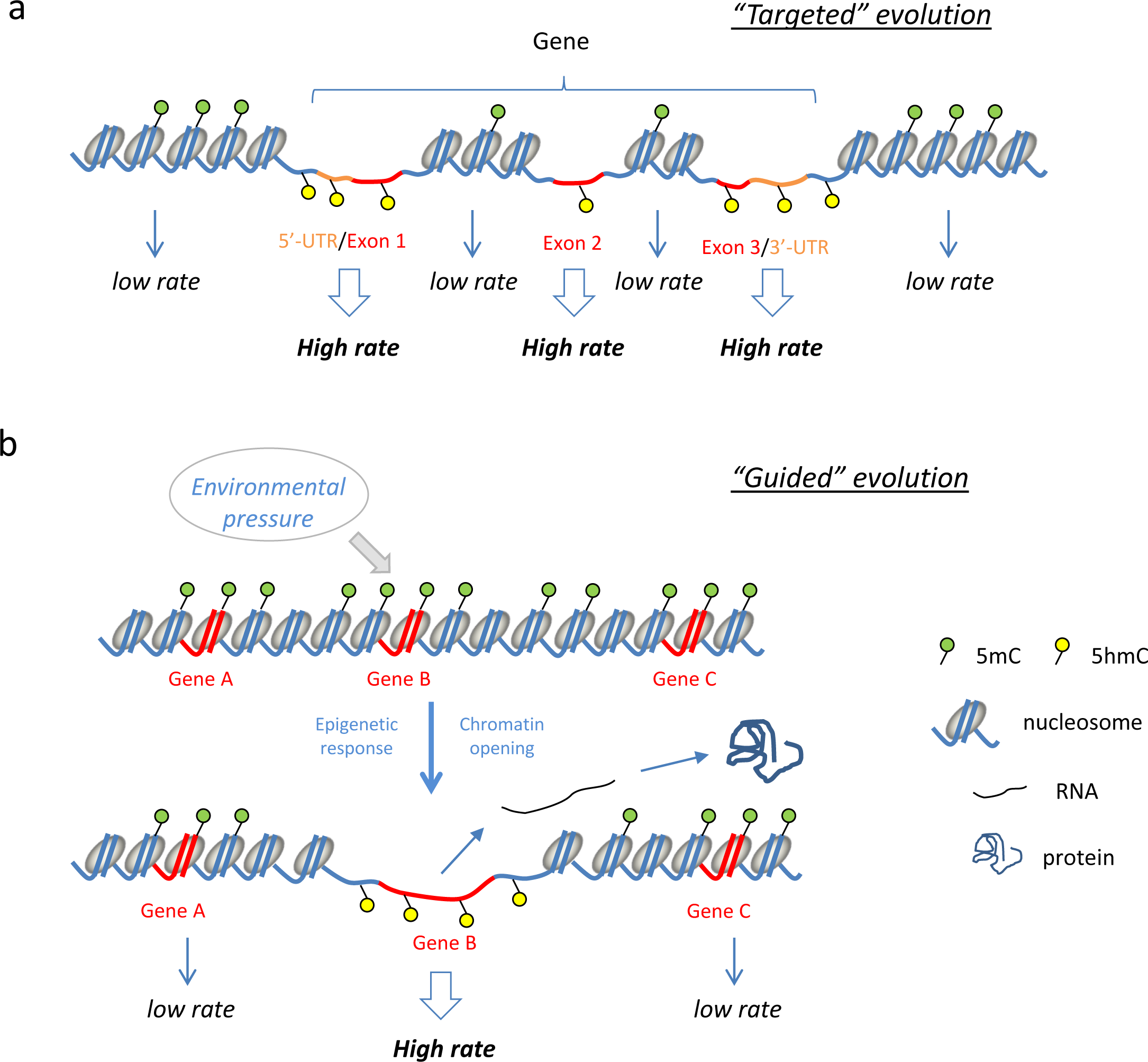
Model of targeted and guided genetic evolution. Based on this study the point recombination is strongly associated with the presence of 5hmC marks. **(a) “Targeted” evolution**. As *bona fide* epigenetic mark, 5hmC, is enriched in functional regions such as UTRs and exons (compare figure 5e). Thus, utilization of this mark by the point-recombination machinery would facilitate a targeted evolution: function-related loci would undergo genetic recombination more frequently than quiescent or non-functional regions. **(b) “Guided” evolution**. 5hmC is also a product of oxidative C-demethylation and therefore indicative of recently opened chromatin. As this process can also occur in response to environmental signals, the newly formed 5hmC marks would direct the recombination machinery also to these sites. In this model, the environmental pressure would therefore “guide” genetic evolution in a Lamarckian sense.

It is already established that ‘Transgenerational epigenetic inheritance’ (TEI) allows the transfer of environmentally induced phenotypes through the germ-line (Rakyan and Whitelaw 2003; Daxinger and Whitelaw 2012; Blake and Watson 2016). Epigenetic imprintment however is only transient and lost after few generations but a link between genetic recombination and TEI may allow to convert “soft” epigenetic marks into “hard” genetic variation. Further experimental studies need to be undertaken to prove the validity of this intriguing hypothesis.

## Methods

### Definition of archaic SNPs

Archaic SNPs are defined here as the set of human SNPs where the derived and ancestral allele is present respectively in the sequence of *denisova hominins* (Meyer et al. 2012a) and chimpanzee (Chimpanzee Sequencing and Analysis Consortium 2005). Human alleles shared with *denisova* were obtained as VCF files (http://cdna.eva.mpg.de/denisova/VCF/hg19_1000g/; (Meyer et al. 2012b)), of which all variants annotated in the 1000 Genome Project (The 1000 Genomes Project Consortium 2010; The 1000 Genomes Project Consortium 2012) were extracted. For each SNP, the ancestral allele is defined to be the one found in the chimpanzee genome (Chimpanzee Sequencing and Analysis Consortium 2005) and the ‘archaic’ derived allele being the one found in the *denisovan* genome. This resulted in a total of 1,897,400 archaic SNPs. All positional information is based on human genome hg19.

### Definition of archaic linkage blocks

The linkage analysis was carried out with phase 3 phased genotype data for 99 individuals of the LWK population (Luhya in Webuye, Kenya) provided by the 1000 genome project (The 1000 Genomes Project Consortium 2010; The 1000 Genomes Project Consortium 2012). The linkage disequilibrium was computed using a Java program, which allowed the computation based on the actual frequency of the derived allele (*f_a_*) instead of just the minor allele frequency. r^2^ values were determined for all possible pairs of derived alleles (*r^2^_a1,a2_*), located within a window of 200,000 bp, using the following equations (with P_a1,a2_ representing the probability of observing the derived alleles a1 and a2 on the same DNA strand):

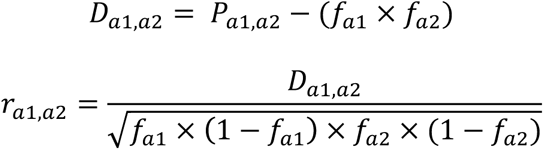

This matrix allowed for the identification of the sets of perfectly linked SNPs (block-SNPs). Their derived alleles represent the preserved core-haplotype of the original archaic haplotype found in *denisovan*. The linkage analysis of derived alleles was carried out in reference to the core-haplotype (r^2^_a,hap_). Archaic linkage blocks were then formed by assigning each of the remaining singleton SNPs of the archaic SNP set (linked SNPs), to the core haplotype with the closest linkage (i.e. highest r^2^_a,hap_ value). This resulted in 237,313 core-haplotypes comprising of 1,021,071 SNPs with 876,193 singleton SNPs in linkage (136 singleton SNPs were omitted as there were no core-haplotypes within 200,000bp).

### Definition of allele frequencies and allelic mobility

Allele frequencies of archaic SNPs of the studied LWK population are expressed as an absolute frequency of the derived allele (f_a_) as well as a normalized frequency (Δf_a,hap_) defined by the difference between the frequency of the derived (f_a_) and the associated core-haplotype (f_hap_). The latter was also used as indicator of expansion or retraction of the derived allele, which was respectively indicated by a positive or negative *Δf_a,hap_* value. For accurate phasing, *Δf_a,hap_* was defined by *f_a_* – *f_hap_* in case of a positive value for r_a_,_hap_, while in case of a negative value *Δf_a,hap_* was defined by (1 - *f_a_*) – *f_hap_*. The LD-based allelic mobility, serving as a proxy for the NCO recombination rate, was defined as 1 - (D’_a,hap_)^2^ (indicated in the text as 1 - D’^2^_a,hap_). D’_a_,_hap_ was computed using the following formula:

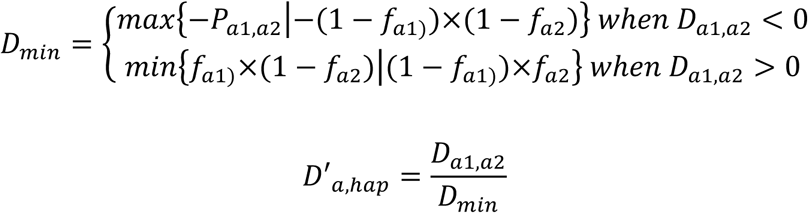

### Assessment of the GC bias

For the analysis of the GC bias only those allele pairs were selected that were characterized by a linkage disequilibrium of D^,2^_a,hap_ = 1 and r^2^_a,hap_ > 1. In this case only 3 allelic combinations existed in the studied population, thereby representing the haplotype sets formed by a single NCO recombination event. Δf_a_,h_ap_ > 0 indicated here the expansion of the derived allele into the ancestral haplotype, while Δf_a_,_hap_ < 0 indicated the replacement of a derived allele by the corresponding ancestral allele:

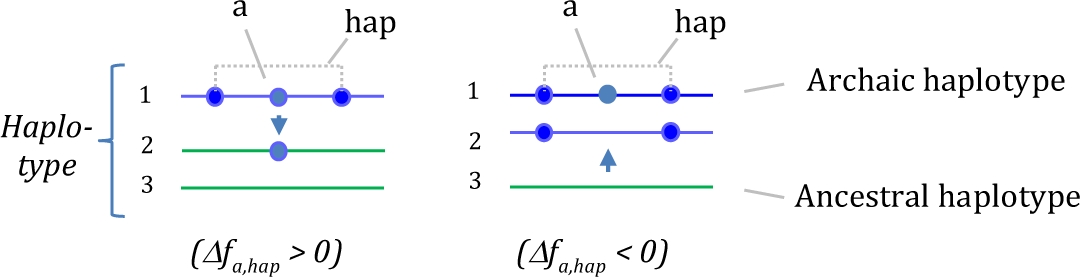

To minimize the influence of genetic drift, only SNPs with a minor allele frequency > 0.2 were considered. Of these, a total of 370,000 non-CpG SNPs and 230,000 CpG SNPs represented heterozygous G:C/A:T pairs. Counting of the SNPs, in which the G:C or the A:T allele was transmitted, revealed respectively 210,000 and 160,000 non-CpG SNPs and 140,000 and 90,000 CpG SNPs, translating into a GC bias of 0.57 (non-CpG SNP) and 0.62 (CpG SNP).

### Frequency of intact ancestral and archaic haplotypes in modern humans

Complete haplotype sets of the 237,312 archaic linkage blocks were generated by downloading the sequence information of the 198 haplotypes of the LWK cohort from the 1000 genome project. The haplotypes, generated by statistical phasing, comprised only ancestral and derived alleles of archaic SNPs. For each block the relative frequency of perfectly preserved archaic haplotypes (entirely consisting of derived alleles) and of perfectly preserved ancestral haplotypes (entirely consisting of ancestral alleles) was determined. The same approach was also used to determine the relative haplotype frequency of mixed haplotypes binned according to the relative fraction of derived alleles (0 < x < 1, increment: 0.1).

### Correlative parameters

Each archaic SNP present in the LWK data set was annotated with a comprehensive set of parameters (details about the data sources are found in Table S1). Frequency of the derived allele (f_a_), normalized allele frequency (Δf_a,hap_) and LD-based allelic mobility (expressed as 1 - D’^2^_a,hap_) were calculated as described above using the genotype data of the east African LWK cohort (99 individuals) (The 1000 Genomes Project Consortium 2010; The 1000 Genomes Project Consortium 2012). Function-related GWAVA (“Genome-wide annotation of variants”) scores (Ritchie et al. 2014) were downloaded together with fitness-related fitCons (“fitness consequences of functional annotation”) scores (Gulko et al. 2015) from the respective sites and assigned to each of the analysed SNPs. Gene annotation, as well as location of genic sub-regions was determined for each SNP using snpEff (PubMed ID 22728672). The average GC-content was determined for each SNP using a window of 200bp (100bp flanking each SNP). Overlap with CpG Islands (CGI) was determined by using data from UCSC Genome Browser (Gardiner-Garden and Frommer 1987; Human Genome Assembly:The Genome Sequencing Consortium 2001; Karolchik et al. 2014). Overlap of the alleles with functionally-defined regions was mostly done with tracks provided by the ENCODE project (Bernstein et al. 2012). This includes tracks of open chromatin (defined by the DNase I tracks) as well as 9 different modifications of histone H3 (K27me3, K4me1, K4me3, K4me2, K9ac, K27ac, K79me2, K36me3, K9me3). For all ENCODE data, an overlap was called when the respective track overlap was detected in any of the tissues and cell types deposited (Bernstein et al. 2012). DMC1 ChIP-Seq tracks generated from human testis (marking meiotic recombination hotspots) were obtained from Pratto *et. al*. (Pratto et al. 2014). Tracks of 5-methylcytosine (5mC) and 5-hydroxymethylcytosine (5hmC) generated from the embryonic stem cell line H1 were obtained from NIH Roadmap Epigenomics (Roadmap Epigenomics Consortium et al. 2015) and Szulwach *et. al*. (Szulwach et al. 2011), respectively.

### Analysis of epigenetically defined sub-regions

111 reference epigenomes divided into 15 epigenetically defined sub-regions were obtained from the NIH Roadmap Epigenomics Consortium (Roadmap Epigenomics Consortium et al. 2015). For each reference genome the average mobility (1 - D’_a,hap_) was computed separately for each sub-region using the LWK-based mobility each SNP had been assigned with. The same procedure was applied to define the state of chromatin (DNase I and 5mC), the presence of 5hmC marks and histone marks (H3K27me3, H3K4me3, H3K36me3 and H3K9me3). Histone marks were defined using the respective tracks provided for each of the reference epigenomes. State of chromatin and 5hmC marks were calculated using the tracks defined for H1 stem cells (Szulwach et al. 2011; Roadmap Epigenomics Consortium et al. 2015). Definition of alleles located proximal or distal to recombination hotspots was based on the overlap with the testis-derived DMC1 tracks obtained from Pratto et al 2014 (Pratto et al. 2014).

### Statistical analysis and data visualization

Data processing and management was done using a combination of Biovia Pipeline Pilot and the R statistical language (version 3.3.1). Statistical analyses were done using the R statistical language (specifics of the statistical tests are described in the respective figure and table legends). Visualization of the data was done using both TIBCO Spotfire and R.

## List of abbreviations

5hmC: 5-hydroxymethylcytosine
5mC: 5-methylcytosine
BivFlank: Bivalent flanking region
bp: Base-pairs
CGI: CpG Islands
ChIP-seq: Chromatin Immunoprecipitation and Sequencing
CO: Crossover
DDR: DNA damage repair
*Denisovan h*: *Denisovan hominid*
DMC1: Meiotic recombination protein, RecA homolog
ENCODE: Encyclopedia of DNA elements
EnhBiv: Bivalent enhancer
f_a_: Absolute frequency of derived alleles
FitCons: Fitness consequence
GWAVA: Genome-wide annotation of variants
*H. sapiens*: *Homo sapiens*
hap: Core haplotype
HLA: Human Leukocyte Antigen
kb: Kilo-basepairs
LD: Linkage disequilibrium
LWK: Luhya from Webuye, Kenya
MHC: Major Histocompatibility Complex
NCO: Non-Crossover
PRC2: Polycomb repressive complex 2
ReprPC: Polycomb Repressed Regions
ReprPCWk: Weak Polycomb Repressed Regions
SNP: Single Nucleotide Polymorphism
TEI: Transgenerational Epigenetic Inheritence
TSS: Transcriptional start site
TssBiv: Bivalent transcriptional start site
Tx: Transcribed Region
TxFlank: Flanking a transcribed region
TxWk: Weakly transcribed region
UTR: Untranslated Region
ZNF: Zinc Finger

## Author’s contributions

B.L. and R.M. created databases and analyzed data, S.C., W.L. and AK.A. provided data and other crucial information, M.P. supervised the computational analysis, O.R. developed the concept and directed the project.

## Acknowledgements

All the Singapore Immunology Network authors are supported by the A*STAR/Singapore Immunology Network core grant and the A*STAR Research Attachment Program (ARAP) for a graduate student involved in the study. We thank A. Torroni, H. Lehrach, M. Hoehe, Y.-A. Lee, and S. Brenner for very helpful discussions.

## Supplemental figures and tables

**SUPPLEMENTAL TABLE 1:** Summary of data sources

| Name | Description | Publication | PMID | URL |
| --- | --- | --- | --- | --- |
| 1000 Genome Project | VCF files with hg19 as the reference were downloaded from the website. | Auton *et. al.* 2015 | 26432245 | http://www.internationalgenome.org/home |
| NIH Roadmap Epigenome | Chromatin state, DNase, histone and methylation data was downloaded from the website. | Kundaje *et. al.* 2015 | 25693563 | http://egg2.wustl.edu/roadmap/web_portal/ |
| GWAVA | BED file containing GWAVA scores were downloaded from the website. | Ritchie *et. al.* 2014 | 24487584 | https://www.sanger.ac.uk/sanger/StatGen_Gwava |
| fitCons | Integrated fitCons scores were downloaded from the website. | Gulko *et. al.* 2015 | 25599402 | http://compgen.cshl.edu/fitCons/ |
| ENCODE | narrowPeak BED files for histone and TF were downloaded from the website. | Dunham *et. al.* 2012 | 22955616 | https://www.encodeproject.org/ |
| ENCODE | DNase data in the form of BED files were downloaded from the UCSC mirror. | Dunham *et. al.* 2012 | 22955616 | https://genome.ucsc.edu/encode/ |
| UCSC | CpG island data was obtained from the track cpgIslandExt for hg19. | Kent *et. al.* 2002 | 12045153 | https://genome.ucsc.edu |
| Recombination hotspot | Data obtained from supplementary data file S1. | Pratto *et. al.* 2014 | 25395542 | https://www.ncbi.nlm.nih.gov/pubmed/25395542 |
| 5hmC marks | Data was obtained from supplementary table S1. | Szulwach *et. al.* 2011 | 21731508 | https://www.ncbi.nlm.nih.gov/pubmed/21731508 |

**SUPPLEMENTAL FIGURE 1:**
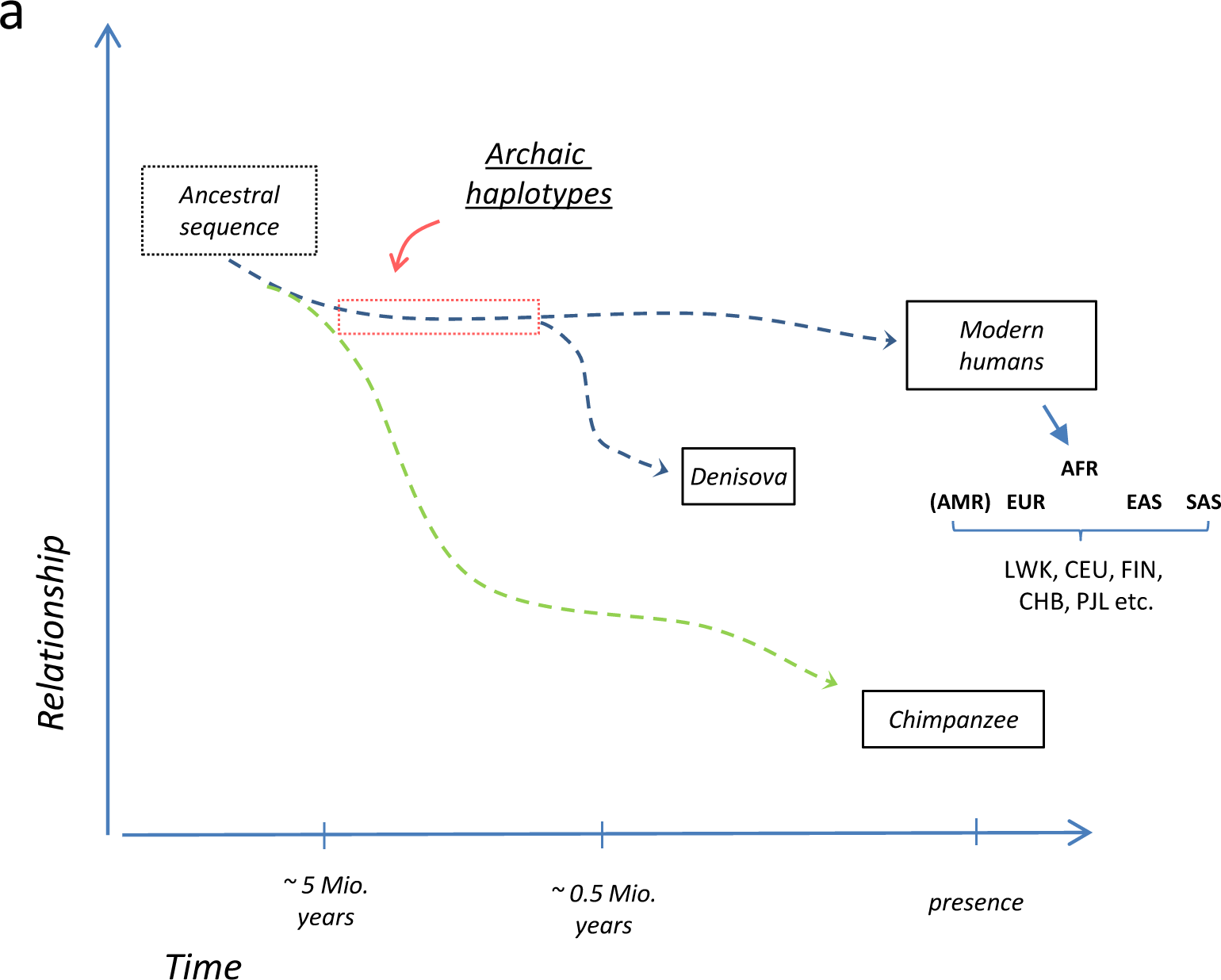
Time lines for the formation of archaic linkage blocks. In this study ‘archaic SNPs’ were defined as SNPs whose derived allele is found in the *denisovan* genomes while the ancestral allele is present in chimpanzee. As a rough estimate, chimpanzee ancestors separated from the *h. sapiens* line about 5 million years ago, while the *denisova h*. lineage split off around 500,000 years ago. This provides a time window of 0.5 – 5 million years, at which point mutations accumulated in the genomes of the common ancestor of *denisova h*. and *h*. sapience to form derived ‘archaic’ haplotypes. After separation, recombination events rearranged the haplotype sequence for about 500,000 years by exchanging derived alleles with the ancestral alleles (and vice versa). Genomic sequence information of *denisova* was obtained from archeologic specimen of individuals, living around 40,000 years ago, while genotype information of modern human was provided by the 1000 Genome Project. AFR: African, AMR: American, EU: European, SAS: South-Asians, East-Asians; LWK: Luhya in Webuye, Kenya (AFR), CEU: Utah Residents with Northern and Western European Ancestry (EUR), FIN: Finnish in Finland (EUR), CHB: Han Chinese in Bejing, China (EAE), PJL: Punjabi from Lahore, Pakistan (SAS).

**SUPPLEMENTAL FIGURE 2:**
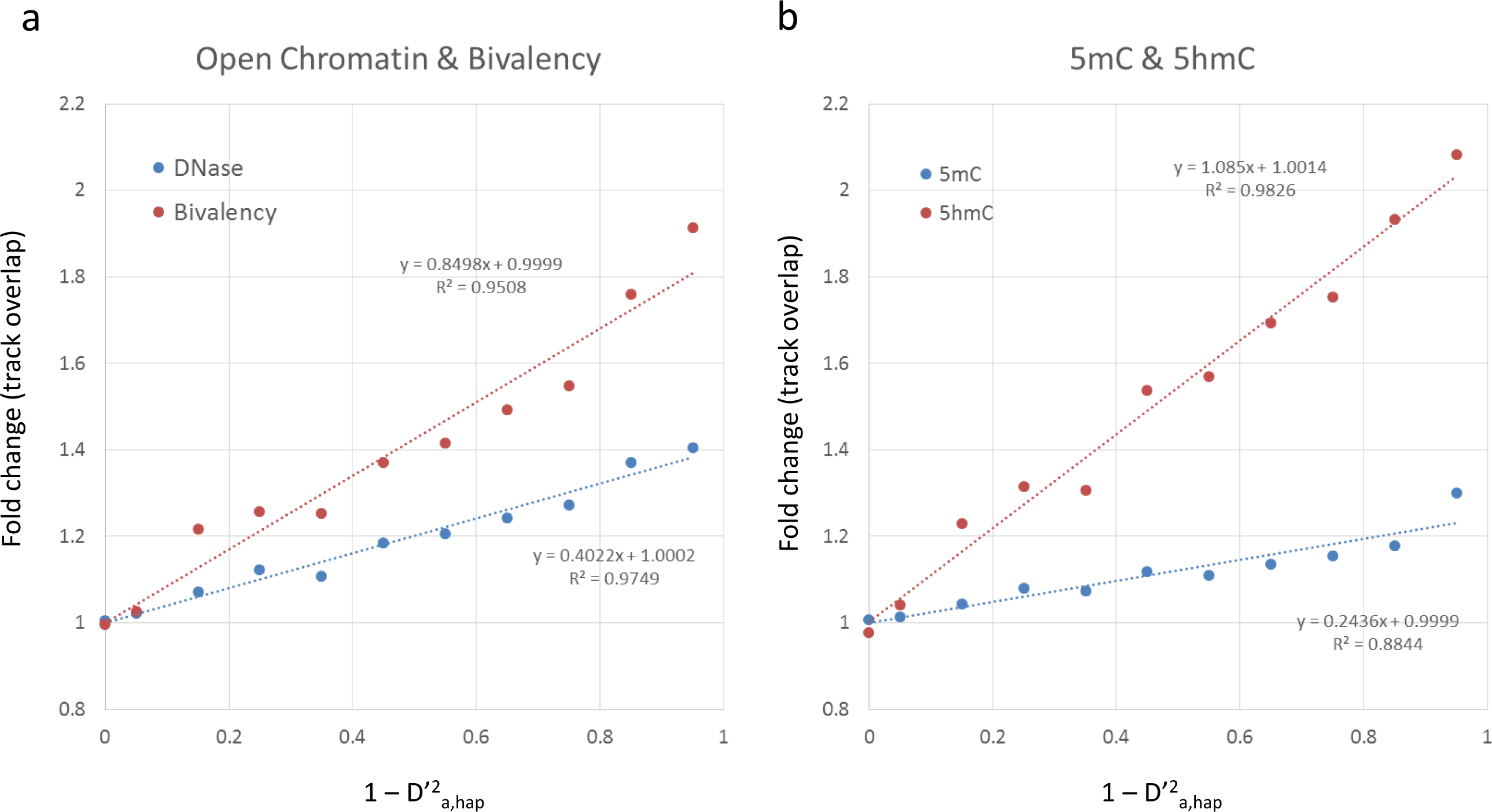
Fold change in open chromatin and epigenetic markers. **(a) Open chromatin and bivalency**. The fold change in the overlap with open chromatin-related DNase I tracks (blue) and H3K27me3/H3K4me3 defined bivalent regions (red) in response to the increase in the average mobility (1 - D’^2^_a,hap_) is shown. The dots indicate the fold change for the binned mobility average (compare figure 4d). Only singleton SNPs are shown. Dashed lines represent linear regressions; R^2^ values and regression parameters are indicated for each curve. Fold change was compiled from the ENCODE data set. **(b) 5mC and 5hmC**. The fold change in the overlap with 5mC tracks (blue) and 5hmC tracks (red) in response to the increase in the average mobility (1 - D’^2^_a,hap_) is shown. The dots indicate the fold change for the binned mobility average (compare figure 5a). Only singleton SNPs are shown. Dashed lines represent linear regressions; R^2^ values and regression parameters are indicated for each curve. Fold change was compiled from data provided for the H1 stem cell.

**SUPPLEMENTAL FIGURE 3:**
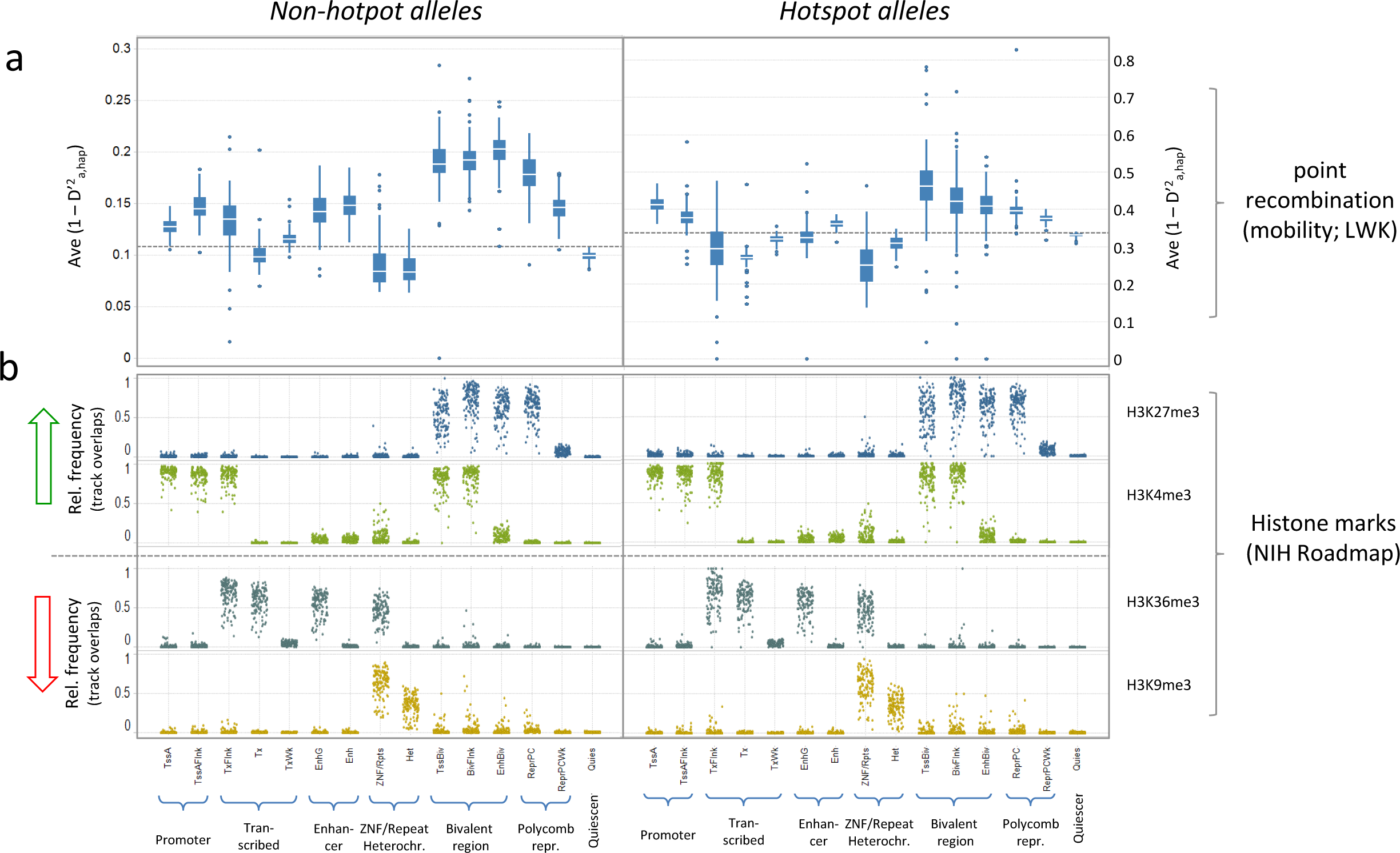
Point recombination and histone marks in epigenetic sub-regions. Point recombination rate and relative frequency histone marks are shown for 15 epigenetic sub-regions defined by the NIH Roadmap Epigenomics Consortium (Roadmap Epigenomics Consortium et al. 2015). TssA: active transcription start site; TssAFlnk: TssA flanking region; Tx: transcribed region; TxWk: weak transcribed region; TxFlnk: Tx flanking region; EnhG: gene-associated enhancer; Enh: enhancer; ZNF: zinc finger nuclear factor; Rpts: repeats; Het: heterochromatin; TssBiv: bivalent transcriptional start site; BivFlnk: bivalent flanking regions; EnhBiv: bivalent enhancer; ReprPC: polycomb repressed region; ReprPCWk: weak polycomb repressed region; Quies: quiescent. The calculations were carried out separately for each of 111 reference genomes using the entire set of 1.9 million archaic SNPs. Separate plots are shown for non-hotspot alleles (left panels) and alleles located at meiotic recombination hotspots (right panels). The assignment is based on their overlap with published DMC1 ChIP-Seq tracks of human testis (Pratto et al. 2014). **(a) Point recombination rate**. The point recombination rate, expressed as average mobility (1 - D’^2^_a,hap_), was calculated using the mobility parameters defined for the LWK cohort (The 1000 Genomes Project Consortium 2010; The 1000 Genomes Project Consortium 2012). The calculation was carried out independently for each reference genomes by using the respective genome-specific region assignments. The boxplots represent the variation of average mobility within these genomes; bars indicate the average mobility of each sub-region. **(b) Histone marks**. The dot plots indicate the local enrichment of the histone marks H3K27me3, H3K4me3, H3K36me3 and H3K9me3. The dots represent the local enrichment of the marks in each of the reference epigenomes. The enrichment is expressed by the relative frequency of archaic alleles overlapping with the tracks (provided by the NIH Roadmap consortium (Roadmap Epigenomics Consortium et al. 2015)) in the respective subregion.

